# Navigating human-dominated landscapes: Spatial strategies of a herd forming ungulate in an agricultural-to-natural matrix

**DOI:** 10.64898/2025.12.02.691804

**Authors:** Ennio Painkow Neto, Gonzalo Barquero, José Manuel Vieira Fragoso

**Affiliations:** Tropical Sustainability Institute (TSI), Carapicuíba, São Paulo, Brazil; Comunidad de Manejo de Fauna Silvestre en la Amazonía y en Latinoamérica (ComFauna), Iquitos, Loreto, Peru; Departamento de Zoologia, Universidade de Brasília, Brasília, Distrito Federal, Brazil; International Union for Conservation of Nature / Peccary Specialist Group (IUCN/SSC), Cambridge, Cambridgeshire, United Kingdom; Instituto Nacional de Pesquisas da Amazônia (INPA/MCTIC), Manaus, Amazonas, Brazil; California Academy of Sciences, San Francisco, California, United States of America

**Keywords:** agriculture, human-wildlife conflict, landscape configuration, movement ecology, *Tayassu pecari*, tropical mosaics, spatial resilience

## Abstract

Landscapes in which preferred, high-quality habitats are embedded can be viewed as a dynamic matrix that can determine the spatial behavior of animal species. In tropical agricultural lands such as Brazil’s Cerrado, understanding how wide-ranging wildlife species navigate these settings is critical for biodiversity conservation. We investigated space-use strategies of white-lipped peccaries (*Tayassu pecari*)—a large-herd forming species and ecological engineer listed as Vulnerable by the IUCN—in a structurally heterogeneous landscape dominated by native vegetation remnants and industrial-scale farms under a legal regulatory framework of agricultural lands, private nature reserves and a national park. Using high-resolution GPS telemetry, we tracked 12 herds across this landscape from 2019 to 2022. Home ranges were estimated using multiple methods (AKDE 95%, KDE 50/95%, MCP 50/95/100%). Landscape composition was classified using MapBiomas land cover data. The herds with the largest ranges (up to 22,166.96 ha) occupied only native vegetation areas, with >99% natural cover, while herds with the smallest ranges (as low as 1,708.72 ha) occurred exclusively in agricultural areas (with >66% anthropogenic cover), and herds in mixed landscapes, composed of agricultural fields intermixed with patches of native vegetation, had ranges between both these extremes. Differences in range size and landscape cover patterns reflect behavioral plasticity in response to matrix configuration, crop availability, and native habitat structure. Our results reveal that landscape configuration and matrix permeability—not habitat amount alone—govern spatial responses in WLPs. Native vegetation patches (Legal Reserves and Permanent Preservation Areas) within crop mosaics functioned as key anchors for herd persistence outside protected areas, reinforcing the idea that such matrices can be conservation assets. Importantly, herds inhabiting agricultural areas exhibited spatial range contraction, which led to conflict with farmers due to their feeding on crops. Increasing peccary occupation in such landscapes is a growing conservation challenge in regions undergoing rapid agricultural expansion. Our work on the movement ecology of a species usually considered as requiring extensive areas of native vegetation indicates its resilience in landscapes dominated by humans. Understanding and managing the landscape matrix is essential for developing coexistence strategies that align biodiversity conservation goals with agricultural realities in the Anthropocene.

## Introduction

The landscape matrix—long viewed as a passive backdrop between habitat fragments—has emerged as a key determinant of ecological, evolutionary, and conservation dynamics in human-modified areas (Fletcher Jr et al., 2024). A growing body of evidence suggests that matrix characteristics can have equal or even greater influence than patch attributes such as area and isolation in determining species persistence, movement patterns, and extinction risk (Prugh et al., 2009; Watling et al., 2011; Ramírez-Delgado et al., 2022). Rather than being considered a uniform non-habitat, the matrix is increasingly recognized as a dynamic and heterogeneous domain that shapes demography, dispersal, gene flow, and selection pressures (Fletcher Jr et al., 2024). This conceptual shift reframes conservation planning: instead of focusing solely on remnant habitat patches, it is now essential to manage the functional quality, configuration, and permeability of the entire landscape matrix, especially in tropical systems undergoing rapid and often irreversible land-use change (Prevedello & Vieira, 2010; Driscoll et al., 2013).

The accelerating transformation of ecosystems due to agriculture, infrastructure expansion, and land-use intensification has triggered a global connectivity crisis, threatening biodiversity and disrupting key ecological processes (Fletcher Jr et al., 2024). Beyond habitat loss alone, it is the reconfiguration of landscapes—and the reduction in functional connectivity—that increasingly constrains animal movement, population viability, and ecosystem resilience (Doherty et al., 2021a; Fletcher Jr et al., 2024).

In response, conservation science has begun to integrate animal movement ecology with landscape ecology to address spatial dynamics in agriculturally dominated landscapes (Doherty & Driscoll, 2018). This integration is crucial to understanding how wide-ranging species navigate mosaics of native vegetation, forest remnants, and croplands. The latter are mostly viewed as anthropogenic barriers (Jorge et al., 2021). Movement ecology, as a conceptual framework, allows researchers to consider the motion and navigation capacities of animal species, and how these interact with landscape features to influence movement decisions and fitness outcomes (Nathan et al., 2008; Allen & Singh, 2016).

A global meta-analysis by Doherty et al. (2021) found that over two-thirds of studied species exhibited movement changes of ≥20% in response to human disturbance. These alterations, ranging from home range expansion to restriction, were shaped by disturbance type, trophic level, body size, and landscape context—highlighting the pervasive effects of anthropogenic pressures across taxa and ecosystems. For example, wild boars (*Sus scrofa*) in European agroecosystems exhibit smaller home ranges in food-rich but structurally simplified landscapes (Fattebert et al., 2017), whereas marsupials in fragmented Australian woodlands expand their spatial use to compensate for resource dispersion (Gardiner et al., 2019).

Yet despite these insights, comparative studies remain scarce for large-bodied, herding herbivores in tropical savannas—species particularly vulnerable due to their extensive spatial requirements, low reproductive rates, and high sensitivity to habitat fragmentation and hunting pressures (Ripple et al., 2015). The Brazilian Cerrado stands as a globally relevant case for investigating wildlife spatial strategies in multifunctional landscapes. As one of the most threatened global biomes, the Cerrado juxtaposes protected areas with expanding agricultural frontiers, remnant private nature reserves (e.g. Legal Reserves), seasonal monocultures, and regenerating habitats—creating a real-world laboratory for applied landscape ecology and conservation planning (De Marco et al., 2023; Pompeu et al., 2024).

In the Cerrado, the white-lipped peccary (*Tayassu pecari*, hereafter WLP) serves as a model species for evaluating space-use responses to landscape modification. WLPs are herd-forming, wide-ranging mammals that rely heavily on resource tracking and social cohesion (Fragoso, 1994, 1998), making them particularly sensitive to habitat loss, fragmentation, and matrix permeability (Reyna-Hurtado et al., 2010; Keuroghlian et al., 2012). Herds move as cohesive units through landscapes (Fragoso, 1998). Ecologically, the WLP is considered an ecosystem engineer and keystone species in Neotropical forests and savannas—playing a pivotal role in nutrient cycling, seed predation, and vegetation dynamics (Fragoso, 2004; Fragoso et al., 2022). In the Cerrado, it is a primary prey species for jaguars (*Panthera onca*) (Foster et al., 2013) and thus important for maintaining trophic interactions and ecological balance in increasingly fragmented landscapes. The WLP is listed as Vulnerable (VU) both globally by the IUCN Red List (Keuroghlian et al., 2013) and nationally in Brazil (MMA, 2022)—reflecting population declines linked to habitat loss, overhunting, and local extinctions across its range (Oshima et al., 2021; Jorge et al., 2021).

Recent research shows that the WLP frequently uses agricultural lands—particularly second crop cornfields—as foraging sites, often intensifying its activity in these areas during the dry season (Jácomo et al., 2013, Painkow Neto et al., 2024). This foraging strategy can generate substantial economic losses for farmers (Lima et al., 2019), but also reveals behavioral flexibility in how the species engages with transformed landscapes (Painkow Neto et al., 2024).

Studies using high-resolution GPS telemetry in the Cerrado-Amazon transition show that the WLP exhibits seasonally variable home range sizes in a mosaic landscape, shaped by land cover composition and crop availability (Costa et al., 2023). These patterns mirror findings from an earlier study in the same region where herds maintained large, stable home ranges in the Cerrado—ranging from 2,700 to over 25,000 ha—and actively incorporate agricultural fields into their core activity areas (Painkow Neto et al., 2024). Together, these findings highlight the importance of viewing the WLP not just as a victim of landscape change, but also as a species capable of adapting to such a mosaic of landscapes by altering its movement to maximize foraging opportunities while relying on forest remnants as refuges.

Here, we investigate how 12 WLP herds navigate an agricultural-to-natural landscape gradient in the Cerrado using high-resolution GPS tracking data. By quantifying home range sizes and habitat use across this mosaic of land uses, our goal is to generate insights into how large mammals respond to landscape heterogeneity under dynamic socio-environmental pressures. Although our empirical focus is on a single Neotropical species, the broader aim is to inform discussions on wildlife management, landscape resilience, and coexistence strategies in human-dominated ecosystems—contributing to the growing body of evidence that emphasizes the prominent role of the matrix in ecology, evolution, and conservation (Iwamura et al., 2014, 2016).

Based on prior studies, we hypothesized that WLPs would face spatial constraints in highly modified landscapes—particularly in intensively cultivated agricultural areas—due to their large spatial requirements, cohesive social structure, and high sensitivity to fragmentation and hunting pressure (Oshima et al., 2021; Jorge et al., 2021; Peterson et al., 2025). However, empirical findings also suggest that in certain agricultural settings, the WLP may reduce its home range size when high-energy crops such as corn are locally abundant and seasonally available (Jácomo et al., 2013; Painkow Neto et al., 2024). We therefore expected that herds would shape their space-use patterns in response to the spatial configuration of native and agricultural habitats, modulated by seasonal crop dynamics and matrix structure. Furthermore, despite sharing a relatively restricted geographic context, we hypothesized that herds could display divergent space-use strategies—driven by fine-scale landscape variation and behavioral flexibility—rather than exhibiting uniform responses to land-use gradients.

## Materials and Methods

### Study Area

This study was conducted in the border region of the States of Goiás, Mato Grosso, and Mato Grosso do Sul, within the southwestern portion of the Cerrado in Brazil (Figure 1). Emas National Park (ENP) served as the central reference area (−18.279982°, −52.900918°; WGS 84 datum), a native vegetation area surrounded by a landscape that has been extensively modified by agricultural expansion (IBAMA, 2004). In addition to formally protected areas such as ENP, Brazil’s Forest Code (Law No. 12,651/2012; Government of Brazil 2012) requires the preservation of native vegetation within private lands through two mechanisms: Permanent Preservation Areas (APPs), which safeguard riparian zones and steep slopes, and Legal Reserves (RLs), which mandate conservation of a proportion of native vegetation. RL requirements vary: 35% in the Cerrado portion of Mato Grosso (within the Legal Amazon) and 20% in Goiás and Mato Grosso do Sul. Since the 1970s, when federal programs such as POLOCENTRO incentivized colonization of the undeveloped “interior”, the region has undergone rapid land-use change (IBDF/FBCN, 1981). Native Cerrado vegetation was initially converted into a mosaic of pastures and croplands; in subsequent decades, pastures were progressively replaced by highly mechanized grain production (Jácomo, 2004). The current matrix is dominated by intensive no-till farming systems, primarily composed of soybeans, corn, cotton, and sugarcane, while open pastures have become rare (MapBiomas, 2023). Farmers typically grow soybeans from October to February, followed by a second crop—usually corn—from February to July. August and September mark the fallow period, during which fields are covered by non-harvested vegetation that serves to protect the soil and enhance nitrogen fixation (Painkow Neto et al., 2024). Crop composition and phenology varied throughout the study period, depending on seasonal and annual planting decisions. Remaining native vegetation—both within ENP and in isolated fragments—is mostly composed of savanna-woodland and native grassland formations typical of the Cerrado, while forest patches and seasonally flooded wetlands are limited in extent (Ribeiro & Walter, 2008; MapBiomas, 2023) (Figure 1).

**Figure 1.**
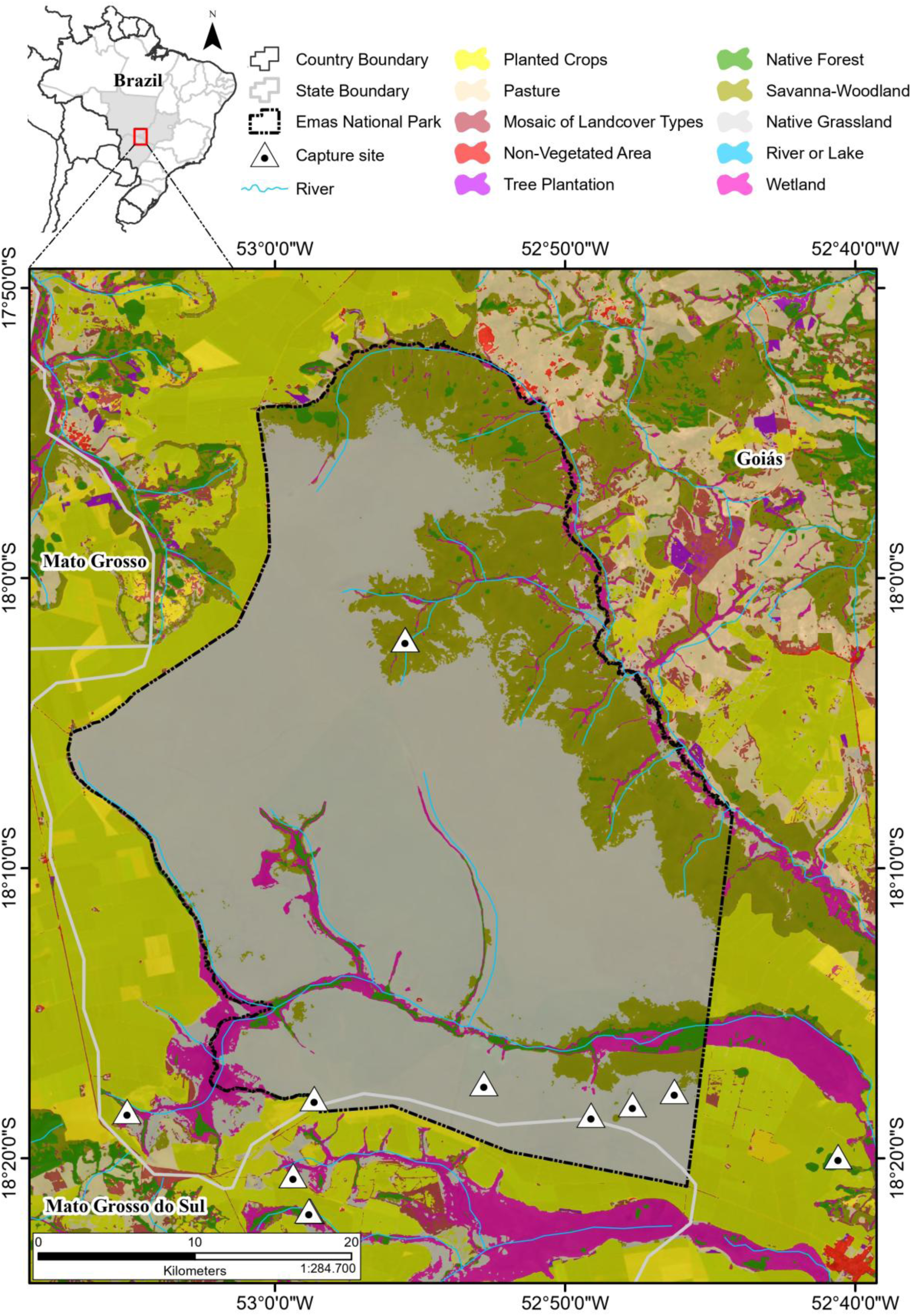
Geographic location of the study area in the Brazilian Cerrado biome, at the intersection of the states of Goiás, Mato Grosso, and Mato Grosso do Sul. Land cover types follow the MapBiomas Project classification (Collection 9; MapBiomas, 2023). White triangle symbols with central black dots indicate GPS collar capture sites of white-lipped peccary (*Tayassu pecari*) herds. The boundary of Emas National Park is shown as a black dashed line. Land cover classes include Planted Crops, Pasture, Mosaic of Landcover Types, Non-Vegetated Area, Tree Plantation (anthropogenic habitats), and Native Forest, Savanna-Woodland, Native Grassland, River or Lake, and Wetland (native habitats).

### Landscape Classification and Habitat Types

To characterize landscape composition and habitat types within each herd’s home range, we used annual land cover maps from the MapBiomas Project – Collection 9.0 (MapBiomas, 2023), selecting the raster layer corresponding to the year in which each herd was tracked (Table S1 in Supporting Information, Figure 1). Raster files were clipped to the spatial extent of the GPS dataset for each herd, based on the specific tracking period.

We reclassified the land cover data into a simplified set of habitat types to distinguish between native and human-modified environments. This functional reclassification was structured as follows:

- Anthropogenic habitats. We retained MapBiomas’ original categories for Pasture and Mosaic of Landcover Types, as they represent transitional or mixed-use areas commonly associated with extensive cattle ranching and rotational land use. All agricultural land-use classes—Soybean, Sugarcane, Cotton, and Other Temporary Crops—were grouped into a single category labeled “Planted Crops,” reflecting the limitations of MapBiomas in capturing within season variability and local-scale accuracy of crop types. Urban Area and Other Non-Vegetated Areas were combined under the category “Non-Vegetated Area”, which includes infrastructure, exposed soil, and human settlements.
- Native habitats. We preserved all natural vegetation classes under the umbrella category “Native Vegetation”, including Native Forest, Savanna-Woodland, Wetland, and Native Grassland —representing the dominant native habitats of the Cerrado biome.

This reclassification provided a clear and consistent framework for comparing habitat composition across home ranges while preserving ecological relevance. For herds tracked over multiple years, we used the land cover map corresponding to the first full year of monitoring as the reference layer, because the agricultural matrix in the region tends to remain relatively unchanged across consecutive years. The resulting habitat maps allowed us to evaluate whether herds primarily occupied native or anthropogenic environments and to assess how landscape composition influenced the extent and configuration of their home ranges.

### Capture and Monitoring of WLPs

Between March 2019 and May 2022, we captured and monitored 12 individual WLPs to estimate home range size and assess habitat use patterns. Corral trap sites were selected based on direct sightings and/or recent signs of activity (e.g., tracks, rooting sites, and resting areas) and knowledge of the location of individual, distinct herds, identified by a specialized field team working continuously in the region since 2017 (Painkow Neto et al., 2024).

To ensure broad representation of landscape contexts and the sampling of distinct herds, capture efforts were strategically distributed across central and southern sectors of Emas National Park, as well as in agricultural areas located south of the park (Figure 1). The range of distances between trapping locations varied considerably, from approximately 100 meters to 11 kilometers. In cases where capture sites were very close to each other (e.g., ∼100 meters), distinct herds had already been identified based on long-term monitoring, and field reconnaissance (Painkow Neto et al., 2024). In the central region of the park, the trap site was located approximately 9.7 kilometers from the park boundary, and the shortest straight-line distance between this central site and traps in the southern region was around 28 kilometers.

To minimize the risk of recapturing individuals from previously monitored herds, site selection was guided by: (i) spatial activity patterns inferred from recent signs and observations; (ii) long-term knowledge of herd distributions within the study area (Painkow Neto et al., 2024); and (iii) field reconnaissance conducted in the days preceding capture events. This approach was essential for capturing variation in space use across contrasting landscapes.

Animals were lured into large corral traps (∼140.4 m²) using corn and salt. Each trap was equipped with an automatic trigger system that closed the entrance gate once animals were inside. Captures were conducted exclusively at night to minimize stress. Once animals were captured, a single adult per herd was selected for collaring based on health and, when possible, social dominance, regardless of sex. Chemical immobilization was performed using Zoletil® 100 (0.07 mL/kg), administered intramuscularly via dart syringe fired from a Teleinject CO₂ rifle. Selected individuals were equipped with GPS collars (Telonics® and Iridium Terrestrial Systems® models). All animals remained in the trap overnight to ensure full recovery of the sedated individual, and the entire herd was released the following morning.

To balance battery life, data storage, and temporal resolution, GPS collars were programmed to acquire locations at either 30-minute or 1-hour intervals. A 30-minute fix rate provides higher temporal resolution, allowing for fine-scale analysis of movement patterns, habitat selection, and behavioral states (Cagnacci et al., 2010; Kays et al., 2015). In contrast, a 1-hour interval extends battery longevity while maintaining sufficient accuracy for robust home range and space-use estimates (Hebblewhite & Haydon, 2010). These settings represent a trade-off between data granularity and deployment duration. Both fix rates are widely used in large mammal studies, particularly for social and wide-ranging species such as WLP. Previous research has shown that fix intervals within this range do not introduce significant bias in home range estimates, regardless of the method applied (Frair et al., 2004; Börger et al., 2006).

### Home Range Estimation and Statistical Analyses

All analyses were conducted in R version 4.3.1 (R Core Team, 2023). GPS data were pre-processed to remove duplicate records, temporal inconsistencies, and spatial outliers, and were reprojected to the SIRGAS 2000 / UTM zone 22S coordinate reference system.

Home ranges were estimated individually for each herd using six complementary methods widely applied in movement ecology, to allow for comparison with past studies: Minimum Convex Polygon (MCP; Mohr, 1947) at 50%, 95%, and 100%; Kernel Density Estimation (KDE; Worton, 1989) at 50% and 95%; and Autocorrelated Kernel Density Estimation (AKDE; Fleming et al., 2015) at 95%. MCP and KDE estimates were calculated using the ‘adehabitatHR’ package (Calenge, 2006), while AKDE estimates were computed using the ‘ctmm’ package (Calabrese et al., 2016), which accounts for temporal autocorrelation in movement data and yields statistically robust estimates of space use.

Landscape composition within home ranges was quantified by spatially intersecting AKDE 95% polygons with annual land cover maps from the MapBiomas Project – Collection 9.0 (MapBiomas, 2023), using the packages ‘sf’ (Pebesma, 2018), ‘terrà (Hijmans, 2023), and ‘raster’ (Hijmans, 2020). For details on raster selection, land cover reclassification, and habitat grouping criteria, see *Landscape Classification and Habitat Types*.

Although all home range estimation methods were compared, AKDE 95% was adopted as the primary metric for interpretation and discussion, in line with recent recommendations in movement ecology for its ability to incorporate spatiotemporal autocorrelation and to produce ecologically meaningful estimates of space use (Fleming et al., 2015).

The distribution of AKDE 95% home range size was assessed using the Shapiro–Wilk test; comparisons between herds were based on the distribution of median values and non-parametric statistical tests were applied if suitable. Differences in home range size and landscape composition among herd groups were tested using the Kruskal–Wallis test. Results are reported using the chi-squared (χ²) test statistic. Post hoc comparisons were conducted using Dunn’s test with Benjamini–Hochberg correction for multiple comparisons. All tests were two-tailed, and statistical significance was set at p ≤ 0.05.

## Results

### Herd Captures and Tracking

Between 25 and 59 individuals were captured per trapping event (Table 2). It was not possible to capture the full herd during the events, and approximately 5 to 20 additional individuals were frequently observed or heard around the corral following a capture. Total herd sizes therefore likely reached 79 individuals, with variation depending on the capture location. Each herd was assigned an acronym-name (Table 2).

**Table 1.**
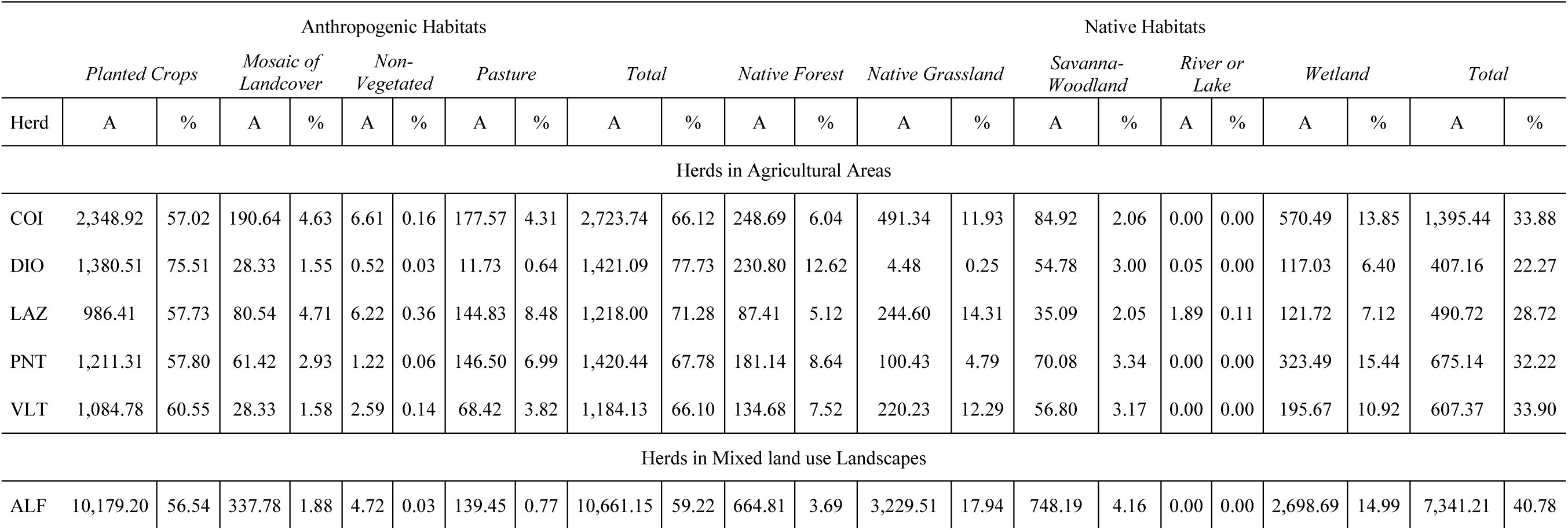
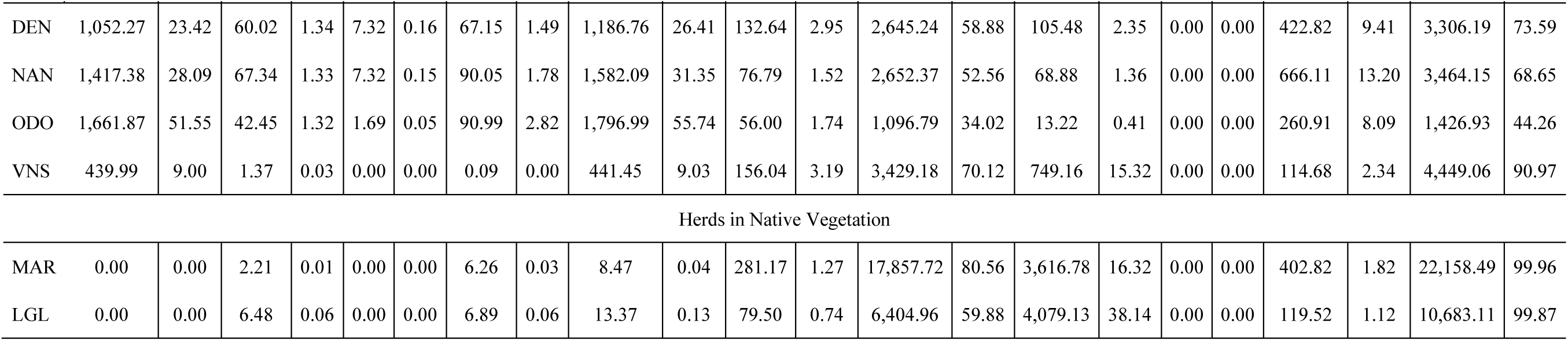
Availability of landscape categories within the AKDE 95% home ranges of 12 white-lipped peccary (*Tayassu pecari*) herds monitored in the Brazilian Cerrado between 2019 and 2022. Herds are organized by herd name and landscape context: Herds in Native Vegetation, Herds in Mixed land use Landscapes, and Herds in Agricultural Areas. Each column labeled “A” indicates the area in hectares (ha) of each land cover category within a herd’s home range; the subsequent column (%) shows the percentage these areas represent relative to the total AKDE 95% home range area. Landscape categories were derived from MapBiomas Collection 9.0 (MapBiomas, 2023) and include Planted Crops, Native Forest, Native Grassland, Mosaic of Landcover Types, Non-Vegetated Area, Pasture, River or Lake, Savanna-Woodland, and Wetland.

**Table 2.** Summary of GPS tracking information for 12 white-lipped peccary (*Tayassu pecari*) herds monitored in the Brazilian Cerrado between 2019 and 2022. Each herd was assigned a name. For each herd, we also report the sex of the collared individual (F = female; M = male), the capture location (PP = Private Property; S-ENP = Southern Emas National Park; C-ENP = Central Emas National Park), the number of individuals captured during the event, capture and tracking dates, duration of monitoring (in days), total number of GPS locations acquired, and the programmed fix rate interval (30 or 60 minutes).

| Herd | Sex | Capture Location | Nº. Individuals Captured | Capture Date | Tracking Start Date | Tracking End Date | Tracking Days | Number of Locations | Fix Rate |
| --- | --- | --- | --- | --- | --- | --- | --- | --- | --- |
| ALF | M | S-ENP | 42 | 06/04/2019 | 14/04/2019 | 12/07/2019 | 90 | 4,185 | 30 |
| COI | M | PP | 33 | 24/03/2021 | 01/04/2021 | 31/12/2021 | 275 | 4,827 | 60 |
| DEN | M | S-ENP | 36 | 03/04/2019 | 11/04/2019 | 25/08/2019 | 136 | 6,264 | 30 |
| DIO | M | PP | 26 | 14/05/2020 | 22/05/2020 | 30/04/2022 | 709 | 15,937 | 60 |
| LAZ | M | PP | 25 | 07/05/2020 | 15/05/2020 | 10/09/2021 | 484 | 11,008 | 60 |
| LGL | F | C-ENP | 35 | 14/07/2021 | 22/07/2021 | 16/05/2022 | 299 | 7,126 | 60 |
| MAR | F | C-ENP | 32 | 23/06/2020 | 01/07/2020 | 11/05/2021 | 315 | 7,464 | 60 |
| NAN | M | S-ENP | 30 | 20/11/2019 | 28/11/2019 | 01/10/2020 | 309 | 14,717 | 30 |
| ODO | F | S-ENP | 59 | 31/03/2019 | 08/04/2019 | 30/08/2019 | 144 | 2,952 | 30 |
| PNT | M | PP | 35 | 03/04/2019 | 12/04/2019 | 25/07/2020 | 471 | 22,212 | 30 |
| VLT | F | PP | 32 | 06/05/2020 | 13/05/2020 | 21/01/2021 | 254 | 4,794 | 60 |
| VNS | F | S-ENP | 31 | 21/08/2019 | 29/08/2019 | 13/01/2020 | 137 | 4,603 | 30 |

**Table 3.** Summary of home range estimates for 12 GPS-collared white-lipped peccary (*Tayassu pecari*) herds monitored between 2019 and 2022 in and around Emas National Park, Brazil. Home ranges were calculated using six methods: minimum convex polygon (MCP) at 50%, 95%, and 100%; fixed kernel density estimation (KDE) at 50% and 95%; and autocorrelated kernel density estimation (AKDE) at 95%. All area values are expressed in hectares. The AKDE 95% estimator was adopted as the primary metric for all subsequent analyses and interpretations throughout the study.

| Herd | AKDE 95% | KDE 50% | KDE 95% | MCP 50% | MCP 95% | MCP 100% |
| --- | --- | --- | --- | --- | --- | --- |
| ALF | 18,002.36 | 574.08 | 7,244.37 | 1,362.84 | 8,807.83 | 12,077.61 |
| COI | 4,119.18 | 822.24 | 3,724.53 | 1,466.72 | 3,341.80 | 4,458.38 |
| DEN | 4,492.94 | 417.25 | 3,052.68 | 787.64 | 3,404.36 | 3,714.94 |
| DIO | 1,828.24 | 279.56 | 1,835.82 | 533.12 | 2,386.48 | 4,070.07 |
| LAZ | 1,708.72 | 151.98 | 1,447.17 | 505.13 | 1,728.17 | 4,697.98 |
| LGL | 10,696.49 | 1,814.82 | 10,563.27 | 2,320.66 | 12,347.30 | 14,829.70 |
| MAR | 22,166.96 | 3,063.97 | 16,532.67 | 6,279.30 | 20,998.65 | 24,513.43 |
| NAN | 5,046.23 | 745.15 | 4,035.80 | 1,087.13 | 5,303.90 | 10,582.91 |
| ODO | 3,223.92 | 219.53 | 1,839.91 | 240.08 | 2,134.63 | 5,012.48 |
| PNT | 2,095.58 | 428.25 | 1,867.90 | 594.18 | 2,114.68 | 4,238.15 |
| VLT | 1,791.50 | 276.36 | 1,807.69 | 659.22 | 2,057.77 | 2,774.62 |
| VNS | 4,890.51 | 979.23 | 4,717.62 | 1,269.61 | 5,141.55 | 6,513.00 |

Landscape contexts had little impact on herd size estimates. In the agricultural areas, herds averaged 30 ± 4 individuals; in the southern region of Emas National Park, around the park boundary, herds averaged 40 ± 12 individuals; and in the central region of the park, herds averaged 34 ± 2 individuals (Table 2).

Tracking durations varied from 90 to 709 days, depending on the fix schedule and logistical conditions (Table 2). This corresponds to a total of 2,952 to 22,212 GPS locations per individual (Figure 2). In the central region of the park, herds were monitored for 299 to 315 days, generating 7,126 to 11,008 locations with a 60-minute fix rate. On agricultural lands, monitoring periods ranged from 254 to 709 days, with 4,794 to 22,212 locations, using 30- or 60-minute fix rate. In the southern boundary region of the park, tracking lasted 90 to 309 days, yielding 2,952 to 14,717 locations, also using 30- or 60-minute fix rate.

**Figure 2.**
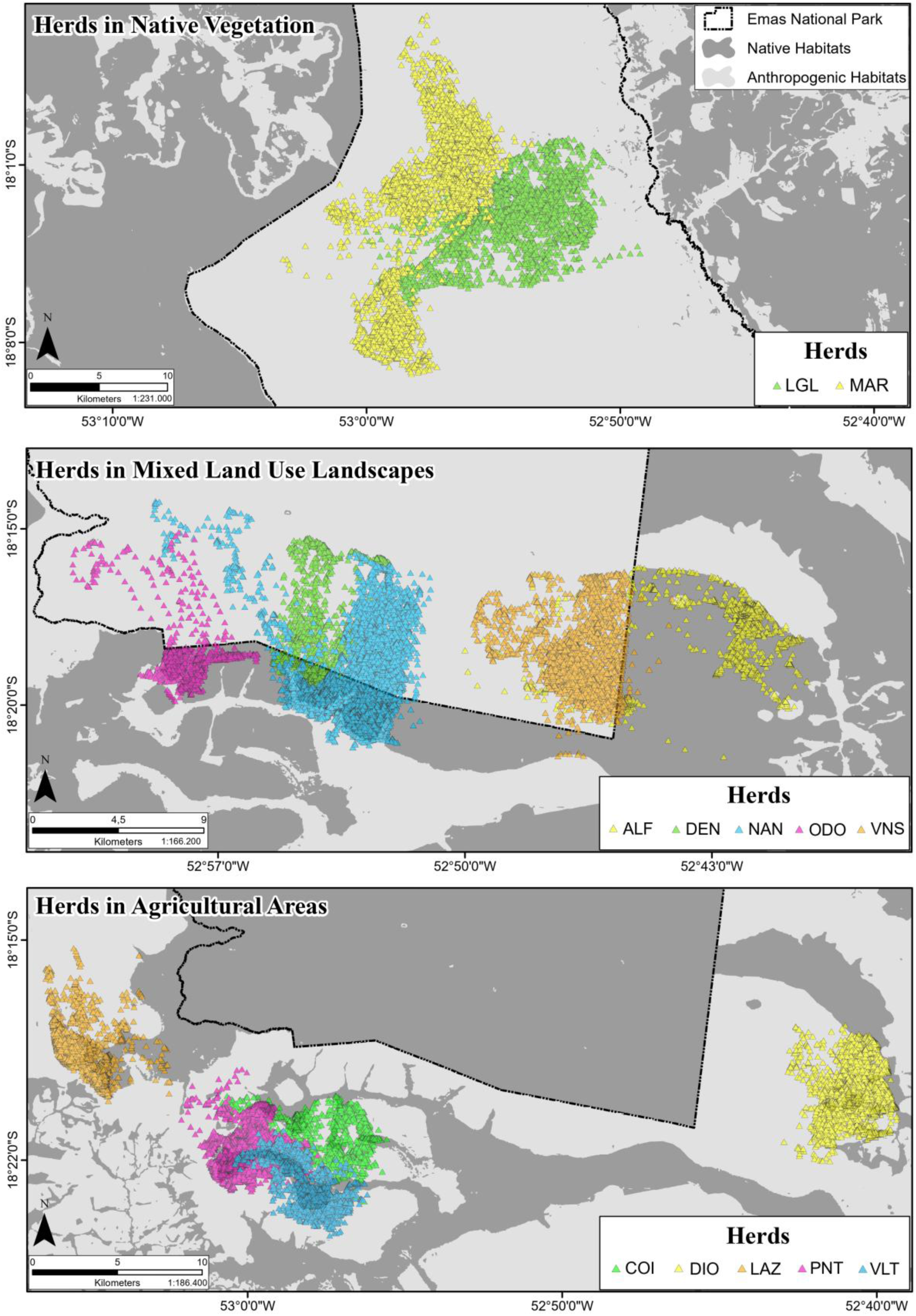
GPS locations of 12 white-lipped peccary (*Tayassu pecari*) herds monitored from 2019 to 2022 in and around Emas National Park, Brazilian Cerrado. Despite minor spatial overlaps visible in the maps, herds remained temporally distinct, with only a single brief co-occurrence observed between two herds in May 2020 (VLT and PNT). Each panel corresponds to a landscape-use category—Agricultural Areas, Mixed land use Landscapes, and Native Vegetation—and displays the GPS locations of herds tracked within each context. Colored symbols represent individual herds. The background land cover was reclassified into two broad habitat types: anthropogenic habitats (including Planted Crops, Pasture, Mosaic of Landcover Types, and Non-Vegetated Area) and native habitats (including Native Forest, Savanna-Woodland, Native Grassland, River or Lake, and Wetland). Light gray areas represent native habitats, dark gray areas represent anthropogenic habitats, illustrating spatial distribution and habitat use across contrasting landscape configurations.

Although plotting of GPS locations reveals that on occasion at home range boundaries animals from one herd used the same space as animals from a neighboring herd (Figure 2), these incursions were highly restricted and the herds did not regularly use the same space at the same time.

### Variation in Home Range Size According to Estimation Method

Home range areas for the 12 collared WLP herds varied among the six methods used. AKDE 95% estimates ranged from 1,708.72 ha (herd LAZ) to 22,166.96 ha (herd MAR). KDE 95% values ranged from 1,447.17 ha (LAZ) to 16,532.67 ha (MAR), while KDE 50% estimates varied between 151.98 ha (LAZ) and 3,063.97 ha (MAR). Using the MCP method, MCP 100% values ranged from 2,774.62 ha (VLT) to 24,513.43 ha (MAR); MCP 95% ranged from 1,728.20 ha (LAZ) to 20,998.65 ha (MAR); and MCP 50% ranged from 240.08 ha (ODO) to 6,279.30 ha (MAR). These differences were consistent across herds and primarily reflect the inherent variation in sensitivity and boundary inclusion among estimation methods (Figure 3).

**Figure 3.**
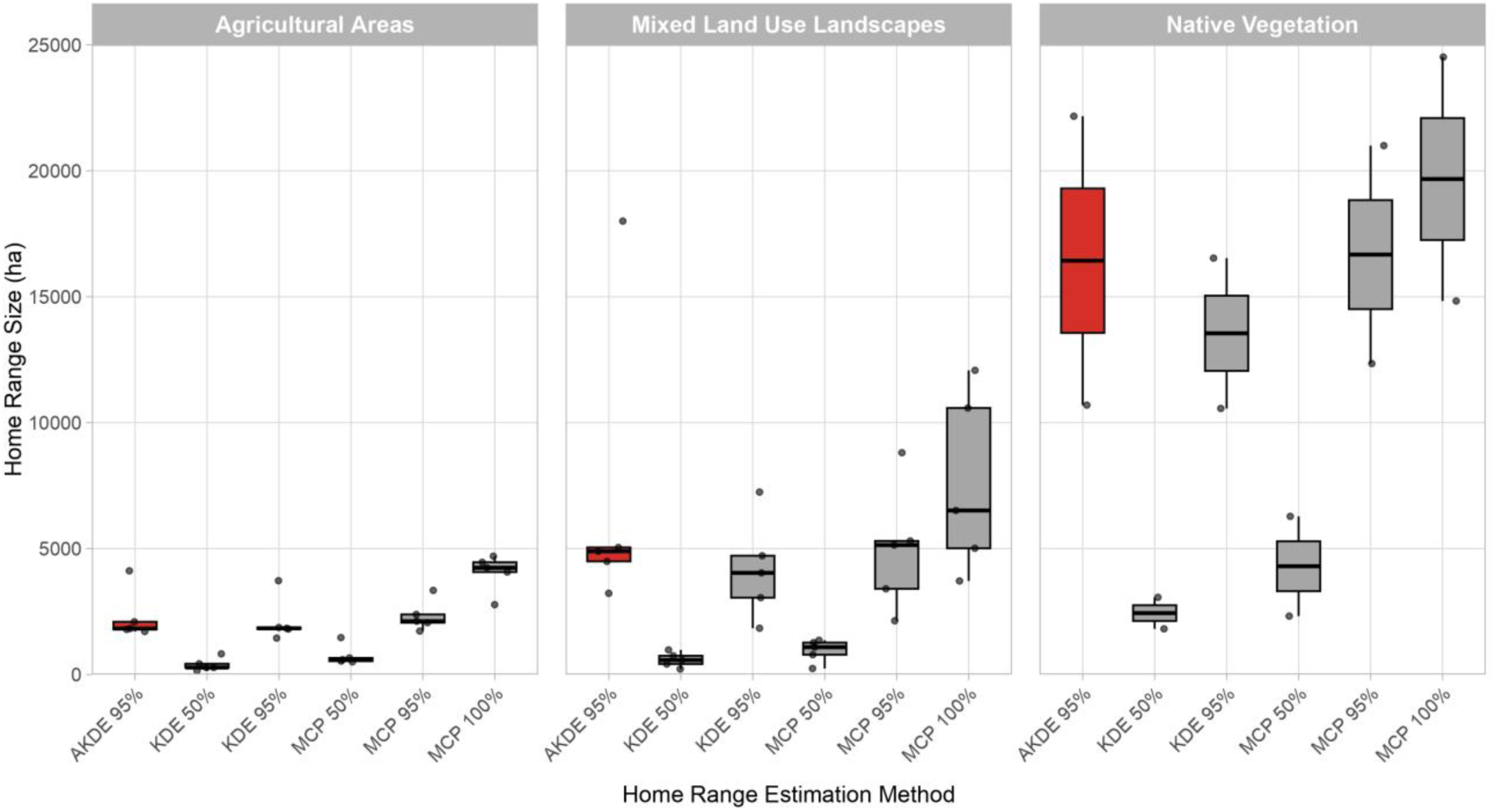
Comparison of home range size estimates (ha) for 12 GPS-collared white-lipped peccary (*Tayassu pecari*) herds monitored in the Brazilian Cerrado, calculated using six analytical methods (AKDE 95%, KDE 50%, KDE 95%, MCP 50%, MCP 95%, and MCP 100%). Herds were grouped into three landscape contexts: Agricultural Areas, Mixed land use Landscapes, and Native Vegetation. Each boxplot represents the distribution of estimates within each group. Boxes display the median and interquartile range (IQR), with whiskers extending to 1.5 × IQR; dots represent individual herd values. Boxplots shown in red highlight AKDE 95%, the estimator adopted as the primary metric for interpretation and discussion in this study. All other methods are shown in gray.

### Herds and Landscape Composition

Analysis of land cover composition within WLP AKDE 95% home ranges revealed three distinct patterns of landscape use: herds in Native Vegetation (MAR, LGL), herds in Mixed Land Use (agricultural fields and adjacent park land) Landscapes (ALF, DEN, NAN, ODO, VNS), and herds in Agricultural Areas (COI, DIO, LAZ, PNT, VLT). This classification emerged from the relative proportions of native and anthropogenic land cover types observed within each home range (*Table 1*).

Across all herds, AKDE 95% home ranges encompassed a wide range of native and anthropogenic land cover classes (*Table 1*). For Native Area herds, native vegetation cover ranged from 99.87% to 99.96%, while anthropogenic cover varied from 0.04% to 0.13% (MAR and LGL, respectively). In Mixed Land Use Landscape herds, anthropogenic cover ranged from 9.03% to 59.22%, and native vegetation from 40.78% to 90.97%. Specifically, the ALF herd presented 59.22% anthropogenic and 40.78% native cover; ODO, 55.74% / 44.26%; NAN, 31.35% / 68.65%; DEN, 26.41% / 73.59%; and VNS, 9.03% / 90.97%. For Agricultural Area herds, anthropogenic land use ranged from 66.10% to 77.73%, while native vegetation ranged from 22.27% to 33.90%. Specifically, the DIO herd exhibited 77.73% anthropogenic and 22.27% native cover; LAZ, 71.28% / 28.72%; PNT, 67.78% / 32.22%; COI, 66.12% / 33.88%; and VLT, 66.10% / 33.90%.

Among native habitats, *Native Grassland* ranged from 0.25% (4.48 ha in DIO) to 80.56% (17,857.72 ha for the MAR herd); *Savanna-Woodland* from 0.41% (13.22 ha in ODO) to 38.14% (4,079.13 ha in LGL); *Native Forest* from 0.74% (79.50 ha in LGL) to 12.62% (230.80 ha in DIO); *Wetland* from 1.12% (119.52 ha in LGL) to 15.44% (323.49 ha in PNT); and *River or Lake* from 0.0029% (0.05 ha in DIO) to 0.11% (1.89 ha in LAZ). Among anthropogenic habitats, *Planted Crops* ranged from 0% (absent in MAR and LGL) to 75.51% (1,380.51 ha in DIO); *Pasture* from 0.09 ha (VNS) to 8.48% (144.83 ha in LAZ); *Mosaic of Landcover Types* from 0.01% (2.21 ha in MAR) to 4.71% (80.54 ha in LAZ); and *Non-Vegetated Areas* from 0.03% (4.72 ha in ALF) to 0.36% (6.22 ha in LAZ).

### Home Range Size and Landscape Composition

We tested whether home range size (AKDE 95%) differed among the three herd categories defined by landscape use (Figure 4). The distribution of AKDE 95% home range size indicated non-normality (Shapiro–Wilk test, W = 0.736, p = 0.0019). The Kruskal–Wallis test revealed a significant difference in home range size across groups (χ² = 8.17, df = 2, p = 0.02). Post hoc comparisons using Dunn’s test with Benjamini–Hochberg correction (Table S2 in Supporting Information) showed that home range sizes of herds in Agricultural Areas were significantly smaller than those of herds in Mixed Land Use landscapes (Z = -2.11, p = 0.05) and in Native Vegetation (Z = -2.59, p = 0.03). Although herds in Mixed Land Use landscapes had smaller home ranges than those in native areas, this difference was not statistically significant (Z = -0.99, p = 0.32).

**Figure 4.**
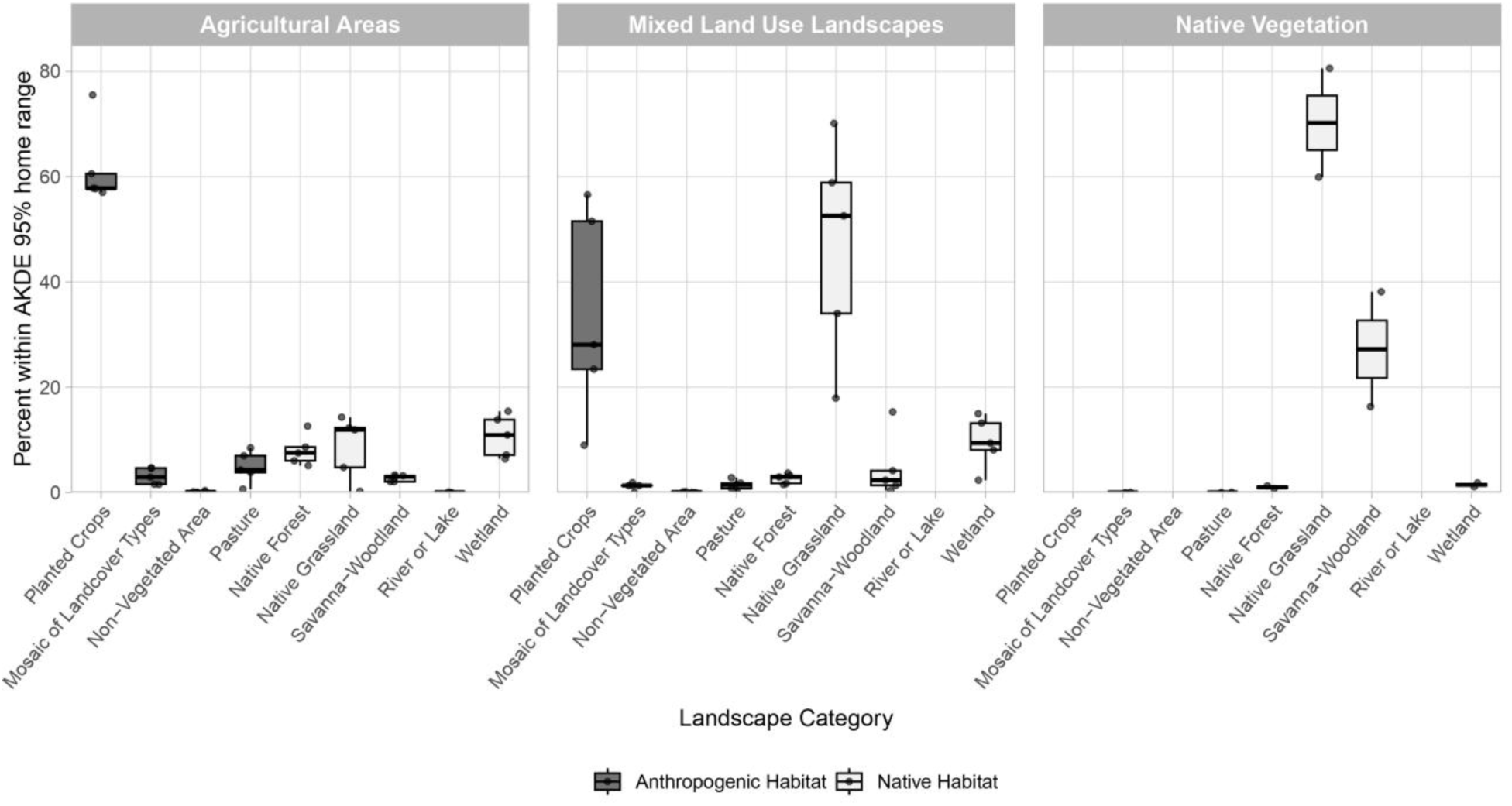
Composition of landscape categories within the AKDE 95% home ranges of 12 GPS-collared white-lipped peccary (*Tayassu pecari*) herds grouped into three landscape contexts: Agricultural Areas, Mixed land use Landscapes, and Native Vegetation. Each facet represents the distribution of habitat types used by herds within a given context. Boxplots show the percentage of each land cover class within home ranges, with boxes indicating the median and interquartile range (IQR), and whiskers extending to 1.5 × IQR; individual herd values are shown as dots. Land cover types were reclassified based on MapBiomas Collection 9.0 into two broad habitat groups: Anthropogenic Habitat (gray 45%) and Native Habitat (gray 95%).

We also assessed whether the percentage composition of landscape categories differed among home ranges of herds in Native Vegetation versus Mixed Land Use, and those in Agricultural Areas (Figure 4). The Kruskal–Wallis test revealed significant differences across groups for Planted Crops (χ² = 9.46, p = 0.01), Native Forest (χ² = 9.42, p = 0.01), Native Grassland (χ² = 8.94, p = 0.01), and Mosaic of Landcover Types (χ² = 7.46, p = 0.02). Dunn’s post hoc test with Benjamini–Hochberg correction (Table S3 in Supporting Information) showed that the proportion of Planted Crops was significantly higher in herds in Agricultural Areas compared to herds in Native Vegetation (Z = 2.82, p = 0.01) and herds in Mixed Land Use landscapes (Z = 2.20, p = 0.04). Similarly, Native Forest was significantly more prevalent in Agricultural Area herds than in Native Vegetation (Z = 2.82, p = 0.01) and Mixed Land Use herds (Z = 2.19, p = 0.04). Native Grassland was significantly less represented in Agricultural Area herds than in Native Vegetation (Z = -2.65, p = 0.02) and Mixed Land Use herds (Z = - 2.28, p = 0.03). Mosaic of Landcover Types was also more prevalent in Agricultural Area herds compared to Native Vegetation herds (Z = 2.52, p = 0.04). Additional comparisons revealed marginal trends: Mosaic of Landcover Types tended to be more represented in Agricultural Area herds than in Mixed Land Use herds (Z = 1.93, p = 0.08); Non-Vegetated Areas were more represented in Agricultural Area herds than in Native Vegetation herds (Z = 2.14, p = 0.10); Pasture was more frequent in Agricultural Area herds than in Native Vegetation herds (Z = 2.22, p = 0.08); Savanna-Woodland was less represented in Agricultural Area herds than in Native Vegetation herds (Z = -1.96, p = 0.08); and Wetlands were more represented in Mixed Land Use herds compared to Native Vegetation herds (Z = 1.89, p = 0.09). No meaningful differences were observed for River or Lake (all p > 0.32), as this class was minimally represented in the areas of all herds, ranging from 0.05 to 1.89 hectares—corresponding to less than 0.11% of the total AKDE 95% home range areas.

## Discussion

### Functional Divergence in Landscape Use

Across the 12 monitored herds, we identified three general patterns of space use based on AKDE 95% home ranges: (i) herds restricted to native habitats in the central core of Emas National Park; (ii) herds limited to intensively cultivated agricultural landscapes interspersed with patches of native vegetation consisting of federally recognized private RLs and APPs; and (iii) herds occupying mixed land use areas that combine protected areas with anthropogenic matrices. These categories reflect differences in spatial arrangement, resource predictability, and habitat connectivity as well as in land cover composition.

Both home range size and habitat composition varied across landscape contexts. WLPs Herds within the national park had access to the highest proportions of natural vegetation (>99.87%) and exhibited the largest home ranges—22,166.96 ha (MAR) and 10,696.49 ha (LGL). These expansive ranges likely reflect the need for broader foraging movements across complex and dynamic resource mosaics. In contrast, the use of areas by herds in intensively farmed regions (e.g., LAZ, DIO, PNT, COI, VLT) contained the highest proportions of anthropogenic cover (66.10–77.73%) and constituted the smallest home ranges, ranging from 1,708.72 to 4,119.18 ha. These results support the hypothesis that spatial contraction occurs in simplified agricultural landscapes, where high-energy, predictable food sources—such as corn—reduce exploratory movement and promote space-use centralization, as described by Morelle et al. (2014) and Painkow Neto et al. (2024).

Herds in mixed land use areas demonstrated intermediate patterns relative to the other groups in both home range size and landscape composition, suggesting flexible behavioral responses to alterations in the landscape matrix, even large-scale change driven by humans. The ALF herd was particularly notable: despite 59.22% anthropogenic cover, it maintained a home range of 18,002.36 ha—the second largest in the study—crossing extensive non-forage sugarcane plantations to access distant corn fields (Painkow Neto et al., 2024). Other herds, such as ODO (3,223.92 ha) and VNS (4,890.51 ha), exhibited more compact ranges. Seasonal dynamics in corn availability likely contributed to this variation; for instance, the monitoring period of VNS coincided with exposed soils and early soybean phenophases, which likely reduced resource availability and habitat attractiveness.

Although spatial overlap occurred among some home ranges, direct intergroup interactions were extremely rare: a sole exception was a brief two-day association between VLT and PNT herds in May 2020, after which both herds resumed independent movements. Even in close proximity, herds maintained spatial exclusivity, reinforcing the species’ strong social cohesion and low tolerance for intergroup overlap—patterns previously described in other studies (e.g., Fragoso, 1998; Reyna-Hurtado et al., 2009).

Collectively, these results reinforce the notion that space use in WLPs is not determined solely by land cover composition, but is strongly modulated by landscape configuration and the functional structure of the overall landscape matrix. The interplay between native vegetation remnants, crop type, and phenology creates a temporally dynamic matrix that shapes the boundaries and scale of space use by WLP. In addition, herd or herd leader memory and experience of travel routes may also play a role, as in movements across non-food sugar cane fields to get to cornfields. Similar foraging movements were observed by Fragoso (1994, 1998) for the species in the Amazon. Thus, there may be a “cultural” or “learned” component to home range patterns. Our findings align with broader theoretical frameworks in movement ecology, which emphasize that configuration and connectivity—rather than habitat amount alone—drive functional space use in fragmented systems (e.g., Prevedello & Vieira, 2010; Doherty & Driscoll, 2018; Fletcher Jr et al., 2024).

### Home Range Variation Across Methods and Contexts

The home range estimates observed in this study—ranging from 1,708.72 ha to 22,167.00 ha using AKDE 95%—are consistent with, and in several cases exceed, those reported for WLP across Neotropical ecosystems (see Painkow Neto et al., 2024). These values reflect both the behavioral and ecological flexibility of the species and the variation in landscape contexts across study sites and methods. The patterns described above in our study are consistent across the different home range estimating methods.

Our findings align with those of Oshima (2019), who reported mean AKDE 95% home ranges of 7,142 ha in the Cerrado and 3,376 ha in the Pantanal, with ranges derived from areas dominated by native vegetation and pasture rather than intensive agriculture. The largest ranges in our dataset—ALF (18,002.36 ha), MAR (22,166.96 ha), and LGL (10,696.49 ha)—were observed in native or mixed land use landscapes, whereas the smallest (e.g., LAZ: 1,708.72 ha) were found in monoculture-dominated agricultural areas, supporting the hypothesis that landscape simplification promotes spatial contraction.

KDE-based estimates from our dataset (1,447.17–16,532.67 ha at 95%) exceeded those reported by Jácomo et al. (2013) in the Cerrado (mean = 7,986.92 ha), which were based on VHF telemetry. The improved spatial and temporal resolution of GPS tracking likely contributed to more precise and ecologically representative estimates of space use. Similarly, Reyna-Hurtado et al. (2009) reported KDE 95% ranges between 3,880 and 9,750 ha in tropical forests of Calakmul, Mexico—values that fall within the range of our mixed and native-context herds (3,052.68 to 16,532.67 ha).

Core area estimates using KDE 50% ranged from 151.98 ha (LAZ) to 3,063.97 ha (MAR), overlapping with those from forested landscapes (e.g., Hofman et al. (2016), who found KDE 50% values between 570 ha and 1,767 ha). These comparisons reinforce the notion that WLPs maintain relatively consistent core activity areas across biomes but adjust total home range size based on resource distribution and landscape structure.

Minimum Convex Polygon (MCP) estimates also demonstrated wide variability. Our MCP 95% estimates ranged from 1,728.17 ha to 20,998.65 ha—overlapping with those reported for Amazonian forests by Fragoso (1998) and the 3,348 ha range documented by Hofman et al., (2016). Notably, the MCP 100% estimate for the MAR herd (24,513.43 ha) greatly exceeded the MCP 100% 2,302 ha range reported by Keuroghlian & Eaton (2008) in the Atlantic Forest, again suggesting that the openness and heterogeneity of the Cerrado may drive broader exploratory behavior.

In agricultural frontiers, Costa et al. (2023) documented MCP 95% home ranges from 1,833 to 7,003 ha, KDE 95% ranges of 2,801 to 7,749 ha, and AKDE 95% ranges of 2,659 to 12,516 ha. These values are generally smaller than those from our native area herds but consistent with those recorded in mixed or agricultural landscapes, supporting the robustness of our findings and the broader applicability of these patterns.

Across methods, a consistent trend emerges: home range size is strongly influenced by landscape context. Larger ranges in native vegetation and mixed land use areas likely reflect greater spatial requirements to access dispersed or temporally variable resources, while smaller ranges in agricultural zones reflect reduced movement due to high-resource density and limited structural complexity. These results confirm that WLPs can thrive even in agricultural areas, as long as some native vegetation remains within the landscape.

### Behavioral Plasticity and Matrix Configuration

WLPs are widely recognized as sensitive indicators of landscape integrity due to their large spatial requirements, cohesive group structure, and heightened vulnerability to fragmentation and hunting (Reyna-Hurtado et al., 2010; Altrichter et al., 2012; Keuroghlian et al., 2015). These traits make them an ideal model for investigating how wide-ranging, large-herd forming mammals respond to environmental heterogeneity in human-modified landscapes. While previous studies have examined WLP space use within distinct biomes—including the Amazon (Fragoso, 1998, 1999), the Pantanal (Keuroghlian et al., 2015; Oshima, 2019), and Mesoamerican forests (Reyna-Hurtado et al., 2009)—few have explored variation in movement behavior at within a single regional mosaic. Our results address this gap, showing that herds monitored within a geographically limited area exhibited strikingly divergent spatial strategies.

This intra-regional variability reinforces the idea that differences in matrix configuration and quality can shape distinct movement responses among conspecifics. This is consistent with findings from other mammals inhabiting mosaics of mixed native and agricultural landscapes (Doherty & Driscoll, 2018; Gardiner et al., 2019), and supports recent advances in landscape ecology that frame the matrix not as a neutral backdrop, but as a key determinant of ecological dynamics (Fletcher Jr et al., 2018, 2024). Our findings also challenge the assumption that high native vegetation cover is a prerequisite for spatial persistence in WLPs. Peterson et al. (2025), for example, modeled movement thresholds in tropical forest landscapes and found that functional connectivity sharply declined below 40% forest cover. While their results underscore the value of native habitat for maintaining connectivity, our data reveal that several herds maintained spatial cohesion, stable home ranges and connectivity even in agricultural landscapes with as little as 22% native cover. These herds occupied environments dominated by intensive agriculture, yet displayed consistent space-use patterns—suggesting a degree of tolerance and resilience to matrix simplification.

Our results do not negate the importance of matrix permeability and connectivity. Rather, they highlight the functional role of certain anthropogenic matrices—particularly those offering predictable, high-energy food sources such as corn—in modulating movement and supporting localized persistence. This behavioral plasticity underscores the need to go beyond habitat amount and consider the structural and functional configuration of the matrix in conservation planning (Prevedello & Vieira, 2010; Driscoll et al., 2013).

### Global Parallels in Movement Ecology of Large Mammals

The spatial responses observed in WLPs mirror patterns increasingly reported for large-bodied mammals across tropical and temperate landscapes. While ecological traits vary widely among species, a recurring trend is evident: the capacity—or necessity—to adjust home range size and spatial strategies in response to landscape structure, matrix composition, and anthropogenic pressures. For example, African forest elephants (*Loxodonta cyclotis*) have been documented persisting in timber extraction zones and agricultural mosaics (Mimeault & Weladji, 2025). However, their home ranges in these modified environments are significantly smaller than in protected areas, suggesting a contraction driven by predictable food availability and increased perceived risk. Similarly, Asian elephants (*Elephas maximus)* in Malaysian agroecosystems adjust their movements according to matrix configuration—displaying smaller ranges and nocturnal foraging behaviors in high-risk, low-cover areas dominated by cropland (Kaliyappan, 2023).

In Europe, wild boars (*Sus scrofa*) exhibit high behavioral plasticity, with spatial patterns strongly shaped by harvest intensity and landscape structure. Fattebert et al. (2017) found that boars inhabiting intensive agricultural zones concentrated their movements in smaller, resource-rich areas, while those in forested or mosaic landscapes maintained larger, more diffuse home ranges. In Australia, macropods such as kangaroos show home range reductions in highly fragmented woodlands, driven by declining connectivity and resource dispersion (Gardiner et al., 2019). Doherty et al. (2021) synthesized evidence from over 200 species globally and demonstrated that human disturbance disrupts animal movement in more than two-thirds of cases, leading to either contraction or expansion depending on local conditions, trophic level, and body size. Such findings highlight the importance of coupling movement ecology with landscape structure to predict spatial responses and inform conservation planning in human-dominated areas.

### Concluding Remarks

Our study demonstrates that land cover, landscape configuration, habitat quality and perhaps most importantly, the landscape matrix along with seasonal dynamics collectively shape space-use strategies in WLP across a heterogeneous land use mosaic. Within a single, limited region of the Brazilian Cerrado, WLP herds exhibited three distinct patterns of landscape occupation—restricted to native vegetation, embedded in intensive agricultural zones, or traversing mixed land use cover types within this matrix. These patterns were not artifacts of geographic distance or biome-level differences but reflected local variation in land cover, including differences in native habitat availability, agricultural crop composition, and matrix permeability.

Our findings reinforce that functional connectivity and matrix structure—not simply habitat amount—drive home range size, movements and spatial behavior in herd forming, wide-ranging species. This aligns with global evidence that large mammals modulate their movements in response to fine-scale environmental change, including that driven by humans, emphasizing the importance of landscape-level conservation planning beyond protected areas. In this context, legal conservation mandates (e.g., RLs and APPs created as part of land use policies in Brazil) and seasonal crop mosaics can enhance ecological functionality when strategically configured to support movement and foraging.

However, the persistence of WLPs in agricultural landscapes faces increasing pressure from the agricultural sector due to their feeding on corn. WLPs can cause substantial crop damage (Lima et al., 2019; Painkow Neto et al., 2024), alongside the lack of governmental wildlife management policies, leading to conflict with producers and instigating control efforts. Proactive engagement of farmers is necessary to prevent local WLP extinction in intensively farmed areas, with cascading effects on ecosystem processes in the remnant native habitats in the landscape mosaic. This scenario underscores the urgent need to align conservation actions with broader societal goals. The global adoption of the United Nations 2030 Agenda for Sustainable Development (UN General Assembly, 2015)—particularly goals related to life on land (SDG 15), zero hunger (SDG 2), and sustainable production (SDG 12)—highlights the importance of multifunctional landscapes that reconcile biodiversity with agricultural productivity. Our findings contribute to this agenda by illustrating that spatial resilience in large mammals is possible even in modified landscapes, provided that matrix elements are managed for ecological function rather than ignored.

## Acknowledgments

We thank CAPES (Coordenação de Aperfeiçoamento de Pessoal de Nível Superior) for providing essential financial support. This research also benefited from contributions by CerradinhoBio, Sindicato Rural de Chapadão do Céu–Goiás, and rural producers located along the southern boundary of Emas National Park. We are especially grateful to the TSI field team—Fernando Marcelino Vicente Lins, Adamo Cardoso Barros, Raphael Francisco Ribeiro, and Mariana Abramo—for their invaluable assistance with data collection. We extend our sincere appreciation to ICMBio, particularly the administration of Emas National Park (ENP), for logistical and institutional support, and to CENAP for technical guidance. We also thank the producers who facilitated field operations, with special recognition to the owners and staff of Fazendas Nova Geração, Agropecuária Duarte, Grupo SUTAL (especially Thomas Peixoto), Grupo Pelizon, Uniggel Sementes, and Fazenda Maraney. Thanks also to Kirsten Silvius for critical reviews and edits. All animal captures and GPS collaring procedures were authorized under SISBIO permit No. 56808 and 69637 issued by the Instituto Chico Mendes de Conservação da Biodiversidade.

## Author Contributions

E. Painkow Neto and G. Barquero conceived the study and conducted fieldwork. E. Painkow Neto curated and analyzed the data and led manuscript writing. G. Barquero contributed to writing and coordinated field operations. J.M.V. Fragoso supervised the study design, provided critical guidance throughout the research process, and contributed to manuscript revisions. All authors approved the final version of the manuscript.

## Conflict of Interest Statement

G. Barquero and E. Painkow Neto received financial and logistical support from Sindicato Rural de Chapadão do Céu–Goiás for work associated with this project. CerradinhoBio and local farmers funded the GPS collars used for peccary tracking. G. Barquero also received consulting support from Sindicato Rural de Chapadão do Céu–Goiás under the project “Projeto Equilíbrio,” established by Technical Cooperation Agreement No. 2/2017–CR-10/CENAP/ICMBio. J.M.V. Fragoso declares no conflicts of interest and received no funding from these entities.

## Notes

### Competing Interest Statement

G. Barquero and E. Painkow Neto received financial and logistical support from Sindicato Rural de Chapadao do Ceu-Goias for work associated with this project. CerradinhoBio and local farmers funded the GPS collars used for peccary tracking. G. Barquero also received consulting support from Sindicato Rural de Chapadao do Ceu-Goias under the project Projeto Equilibrio, established by Technical Cooperation Agreement No. 2/2017-CR-10/CENAP/ICMBio. J.M.V. Fragoso declares no conflicts of interest and received no funding from these entities.

