## Supporting Information for "Navigating human-dominated landscapes: Spatial strategies of a herd forming ungulate in an agricultural-to-natural matrix"

Table S1. Description of land cover categories used in this study, based on the MapBiomas Collection 9 classification system. Land cover classes were reclassified and adapted from the original MapBiomas Collection 9 dataset (MapBiomas, 2023), which provides annual land use and land cover maps for Brazil at 30-meter resolution. Each class was assigned an internationally standardized label to ensure consistency and clarity in ecological analyses and facilitate comparisons with global datasets. Descriptions reflect the biophysical characteristics and dominant land uses associated with each class, including distinctions between planted and natural vegetation, water bodies, anthropogenic mosaics, and savanna formations typical of the Brazilian Cerrado.

| Land Cover Category | Description |
|---|---|
| Planted Crops | Areas used for cultivated crops, including monocultures of soybean, sugarcane, corn, cotton, as well as temporary or perennial crops. |
| Mosaic of Landcover Types | Agropastoral land where it was not possible to distinguish between cropland and pasture. May include abandoned pasturelands under early stages of native vegetation regrowth, anthropized areas within protected zones (excluding Environmental Protection Areas and Indigenous Lands), and peri-urban zones such as smallholdings, rural properties, and residential sites. |
| Non-Vegetated Area | Non-vegetated surfaces including areas with high density of buildings and roads, urban infrastructure, bare soil due to erosion or off-season croplands, and mining areas (industrial or artisanal). Includes regions confirmed by spatial references from CPRM (CPRM, 2023), AHK (Brasilien, 2022), DETER (INPE, 2023) and Instituto Socioambiental (ISA 2022). |
| Pasture | Planted pastures directly related to agricultural activities. Natural pastures are classified as native grasslands or seasonally flooded grasslands and may or may not be subjected to grazing. |
| Native Forest | Vegetation types dominated by native tree species forming a continuous canopy, including riparian forests, gallery forests, dry forests, and cerradão, as well as semideciduous seasonal forests (Ribeiro and Walter 2008). |
| Native Grassland | Open vegetation formations dominated by herbaceous layers, such as clean grasslands, shrubby grasslands, and rocky outcrop grasslands. Some areas may include savanna-like elements such as rocky savanna. |
| Savanna-Woodland | Savanna formations with well-defined herbaceous, shrub, and tree strata, including dense cerrado, typical cerrado, sparse cerrado, and rocky cerrado formations. |

| Land Cover Category | Description |
| --- | --- |
| River or Lake | Rivers, lakes, water reservoirs, and other freshwater bodies, including artificial lakes used for aquaculture or salt production. |
| Wetland | Herbaceous-dominated vegetation subject to seasonal flooding (e.g., wet grasslands) or influenced by river or lake dynamics (e.g., marshes). In some regions, the herbaceous matrix occurs with savanna-type trees or palm formations. |

12

Table S2. Pairwise comparisons of AKDE 95% home range size among herd groups — Herds in Native Vegetation, Herds in Mixed land use Landscapes, and Herds in Agricultural Areas — based on Dunn's post hoc test with Benjamini–Hochberg correction, following a Kruskal–Wallis rank-sum test. Columns show the pairwise comparison, Z statistic, unadjusted *p*-value, and *p*-value adjusted for multiple comparisons. Asterisks (*) denote statistically significant differences based on adjusted p-values ($p \leq 0.05$).

| Comparison | Z value | Unadjusted *p*-value | Adjusted *p*-value |
|---|---|---|---|
| Herds in Agricultural Areas - Herds in Mixed land use Landscapes* | -2.11 | 0.03 | 0.05 |
| Herds in Agricultural Areas - Herds in Native Vegetation* | -2.59 | 0.01 | 0.03 |
| Herds in Mixed land use Landscapes - Herds in Native Vegetation | -0.99 | 0.32 | 0.32 |

Table S3. Pairwise comparisons of landscape category composition among herd groups — Herds in Native Vegetation, Herds in Mixed land use Landscapes, and Herds in Agricultural Areas — based on Dunn's post hoc test with Benjamini–Hochberg correction, following a Kruskal–Wallis rank-sum test. Columns display the landscape category, pairwise group comparison, Z statistic, unadjusted p-value, and p-value adjusted for multiple comparisons. Asterisks (*) indicate statistically significant differences based on adjusted p-values ($p \leq 0.05$). All landscape categories are included, regardless of significance.

| Landscape Category | Comparison | Z value | Unadjusted *p*-value | Adjusted *p*-value |
|---|---|---|---|---|
| Planted Crops | Herds in Agricultural Areas - Herds in Native Vegetation* | 2.82 | 0.00 | 0.01 |
| | Herds in Agricultural Areas - Herds in Mixed land use Landscapes* | 2.20 | 0.03 | 0.04 |
| | Herds in Mixed land use Landscapes - Herds in Native Vegetation | 1.16 | 0.25 | 0.25 |
| Native Forest | Herds in Agricultural Areas - Herds in Native Vegetation* | 2.82 | 0.00 | 0.01 |
| | Herds in Agricultural Areas - Herds in Mixed land use Landscapes* | 2.19 | 0.03 | 0.04 |
| | Herds in Mixed land use Landscapes - Herds in Native Vegetation | 1.16 | 0.25 | 0.25 |
| Native Grassland | Herds in Agricultural Areas - Herds in Native Vegetation* | -2.65 | 0.01 | 0.02 |
| | Herds in Agricultural Areas - Herds in Mixed land use Landscapes* | -2.28 | 0.02 | 0.03 |
| | Herds in Mixed land use Landscapes - Herds in Native Vegetation | -0.93 | 0.35 | 0.35 |
| Mosaic of Landcover Types | Herds in Agricultural Areas - Herds in Native Vegetation* | 2.52 | 0.01 | 0.04 |
| | Herds in Agricultural Areas - Herds in Mixed land use Landscapes | 1.93 | 0.05 | 0.08 |
| | Herds in Mixed land use Landscapes - Herds in Native Vegetation | 1.06 | 0.29 | 0.29 |
| Non-Vegetated Area | Herds in Agricultural Areas - Herds in Native Vegetation | 2.14 | 0.03 | 0.10 |
| | Herds in Mixed land use Landscapes - Herds in Native Vegetation | 1.47 | 0.14 | 0.21 |
| | Herds in Agricultural Areas - Herds in Mixed Land Use Landscapes | 0.88 | 0.38 | 0.38 |
| Pasture | Herds in Agricultural Areas - Herds in Native Vegetation | 2.22 | 0.03 | 0.08 |
| | Herds in Agricultural Areas - Herds in Mixed Land Use Landscapes | 1.67 | 0.10 | 0.14 |
| | Herds in Mixed Land Use Landscapes - Herds in Native Vegetation | 0.96 | 0.34 | 0.34 |
| River or Lake | Herds in Agricultural Areas - Herds in Mixed Land Use Landscapes | 1.62 | 0.11 | 0.32 |
| | Herds in Agricultural Areas - Herds in Native Vegetation | 1.22 | 0.22 | 0.33 |
| | Herds in Mixed Land Use Landscapes - Herds in Native Vegetation | 0.00 | 1.00 | 1.00 |

| Landscape Category | Comparison | Z value | Unadjusted *p*-value | Adjusted *p*-value |
|---|---|---|---|---|
| Savanna-Woodland | Herds in Agricultural Areas - Herds in Native Vegetation | -1.96 | 0.05 | 0.08 |
| | Herds in Mixed Land Use Landscapes - Herds in Native Vegetation | -2.02 | 0.04 | 0.13 |
| | Herds in Agricultural Areas - Herds in Mixed Land Use Landscapes | 0.09 | 0.93 | 0.93 |
| Wetland | Herds in Mixed Land Use Landscapes - Herds in Native Vegetation | 1.89 | 0.06 | 0.09 |
| | Herds in Agricultural Areas - Herds in Native Vegetation | 2.09 | 0.04 | 0.11 |
| | Herds in Agricultural Areas - Herds in Mixed Land Use Landscapes | 0.26 | 0.79 | 0.79 |
